# Dynamic fluctuations of the locus coeruleus-norepinephrine system underlie sleep state transitions

**DOI:** 10.1101/2020.09.01.274977

**Authors:** Celia Kjaerby, Mie Andersen, Natalie Hauglund, Fengfei Ding, Wei Wang, Qiwu Xu, Saiyue Deng, Ning Kang, Sisi Peng, Qian Sun, Camilla Dall, Peter Kusk Jørgensen, Jiesi Feng, Yulong Li, Pia Weikop, Hajime Hirase, Maiken Nedergaard

## Abstract

We normally regard sleep and wake as two distinct opposing brain states, where sleep requires silence of wake-promoting structures such as the locus coeruleus (LC)-norepinephrine (NE) system. We set out to investigate how cortical NE dynamics and NE-related astrocytic activity relates to LC population activity during sleep states.

We show that LC displays regular phasic activity bouts during NREM sleep leading to a slow oscillatory pattern of prefrontal NE levels of which the majority of NE increases does not lead to awakening. NE troughs link to sleep spindles and continued NE decline transitions into REM sleep. Last, we show that prefrontal astrocytes have reduced sensitivity towards NE during sleep.

Our results suggest that dynamic changes in the activity of wake-promoting systems during sleep create alternation between crucial sleep processes and broadening of sensitivity towards incoming sensory input.

**Highlights:**

- Extracellular levels of norepinephrine display dynamic changes during NREM and REM sleep
- Phasic activity of locus coeruleus neurons during NREM underlies slow norepinephrine oscillations
- Spindles occur at norepinephrine troughs and are abolished by norepinephrine increases
- Increased spindles prior to REM reflect the beginning of a long-lasting norepinephrine decline
- REM episodes are characterized by a sub-threshold continuous norepinephrine decline
- The responsiveness of astrocytic Ca^2+^ to norepinephrine is reduced during sleep

## Introduction

Shifts in brain states are needed in order to continuously adapt to changing external and internal conditions. Awake attentive states are fundamental for adaptive behavioral responses, yet equally important is the ability to suppress wakefulness in order to satisfy homeostatic sleep demands (Aston-Jones and Cohen, 2005; Brown et al., 2012).

The locus coeruleus (LC) is considered intimately linked to wakefulness. LC is a small brain stem nucleus with widespread projections, and it is the sole source of norepinephrine (NE) producing neurons in the brain. States of wakefulness and attention coincide with bursting patterns of LC (Aston-Jones and Bloom, 1981), and optogenetic and DREADD-induced activation of LC promotes wakefulness (Carter et al., 2010; Hayat et al., 2020; Li et al., 2016; Porter-Stransky et al., 2019). The conventional view is that LC neurons during sleep is quiescent in comparison to the awake state (Hobson et al., 1975). However, this only holds true for REM sleep (Aston-Jones and Bloom, 1981; Rasmussen et al., 1986); LC displays tonic firing during slow-oscillatory NREM sleep in rats and monkeys (Aston-Jones and Bloom, 1981; Eschenko and Sara, 2008; Eschenko et al., 2012; Foote et al., 1980; Rasmussen et al., 1986).

Previous studies suggest that LC activity during sleep might shape sleep stage transitions. Sleep spindles are 1-2 sec sigma oscillations (8-15 Hz) most frequently observed in cortical regions during NREM sleep stage 2, while they are absent during NREM sleep stage 3, where delta waves dominate (Andrillon et al., 2011). In monkeys and rats, spindle activity coincides with episodes of LC silence (Aston-Jones and Bloom, 1981; Rajkowski et al., 1994; Swift et al., 2018), while LC spiking terminates spindles (Swift et al., 2018). Furthermore, sustained low-frequency optogenetic activation of LC during sleep suppresses spindle occurrences (Hayat et al., 2020; Swift et al., 2018).

The downstream effect of LC activation is release of NE. Cortical extracellular levels of neuromodulators including NE regulates transitions between sleep and wake states (Ding et al., 2016). The density of LC terminals and the level of norepinephrine release are greater in medial prefrontal cortex (mPFC) compared to other cortical areas (Agster et al., 2013; Bellesi et al., 2016; Chandler et al., 2014). Microdialysis studies in mPFC and subcortical regions show that the extracellular level of NE is highest during wake, lower during NREM and lowest during REM (Léna et al., 2005; Shouse et al., 2000). However, due to the temporal constrains of microdialysis, the real-time change in extracellular cortical NE levels in response to LC activity is not known.

Recent evidence suggests that astrocytes may be an integral part of the LC-NE response. Astrocytes display brain-wide increases in Ca^2+^ activity following arousal through LC-NE mediated activation of α1 noradrenergic receptors (Bekar et al., 2008; Ding et al., 2013; Paukert et al., 2014). Also, astrocytic Ca^2+^ responses are absent during sleep and linked to awakenings (Bojarskaite et al., 2020) and DREADD-mediated activation of astrocytic Ca^2+^ signaling increases the latency to fall asleep (Porter-Stransky et al., 2019). However, astrocytes might also be a requisite for normal sleep architecture. Compromising astrocytic Ca^2+^ signaling leads to more fragmented sleep and increased number of spindles (Bojarskaite et al., 2020) and astrocytic Ca^2+^ responses are reduced in response to sedative and anticonvulsant treatment (Thrane et al., 2012; Tian et al., 2005).

While these studies lend support for LC-NE mediated regulation of sleep states, it is still unknown how LC activity affects real-time prefrontal NE levels. Are LC activity and prefrontal NE separate in time? How rapid are NE cleared when LC goes silent? Does the responsiveness of astrocytes to LC-NE depend on brain state? In this study, we took advantage of a newly developed fluorescent NE biosensor (Feng et al., 2019) to correlate prefrontal NE levels with LC activity and prefrontal astrocytic Ca^2+^ responses during states of quiet wakefulness and sleep in mice. Our results suggest that progressive reductions in the activity of wake-promoting systems such as LC-NE and astrocytes is an integral and dynamic part of sleep processes and transitions.

## Results

### Prefrontal NE levels show dynamic fluctuation during NREM sleep

Using TH-Cre driver mice, we expressed GCaMP6f in LC neurons and the newly developed NE biosensor GRAB_NE2m_ (Feng et al., 2019) in mPFC neurons (Figure 1A, Suppl. Figure S1 and S2). We performed simultaneous measurements of LC activity and extracellular NE levels in mPFC using fiber photometry combined with electroencephalography (EEG) and electromyography (EMG) during natural sleep (Figures 1A-B). Based on EEG and EMG recordings, we classified brain states into awake, rapid eye movement (REM) sleep, and non-REM (NREM) sleep. Further, we categorized wake bouts of < 5 s as microarousals (Figure 1B). In a subset of animals, sleep-to-wake transitions were analyzed based on mobility (Suppl. Figure S3).

**Figure 1.**
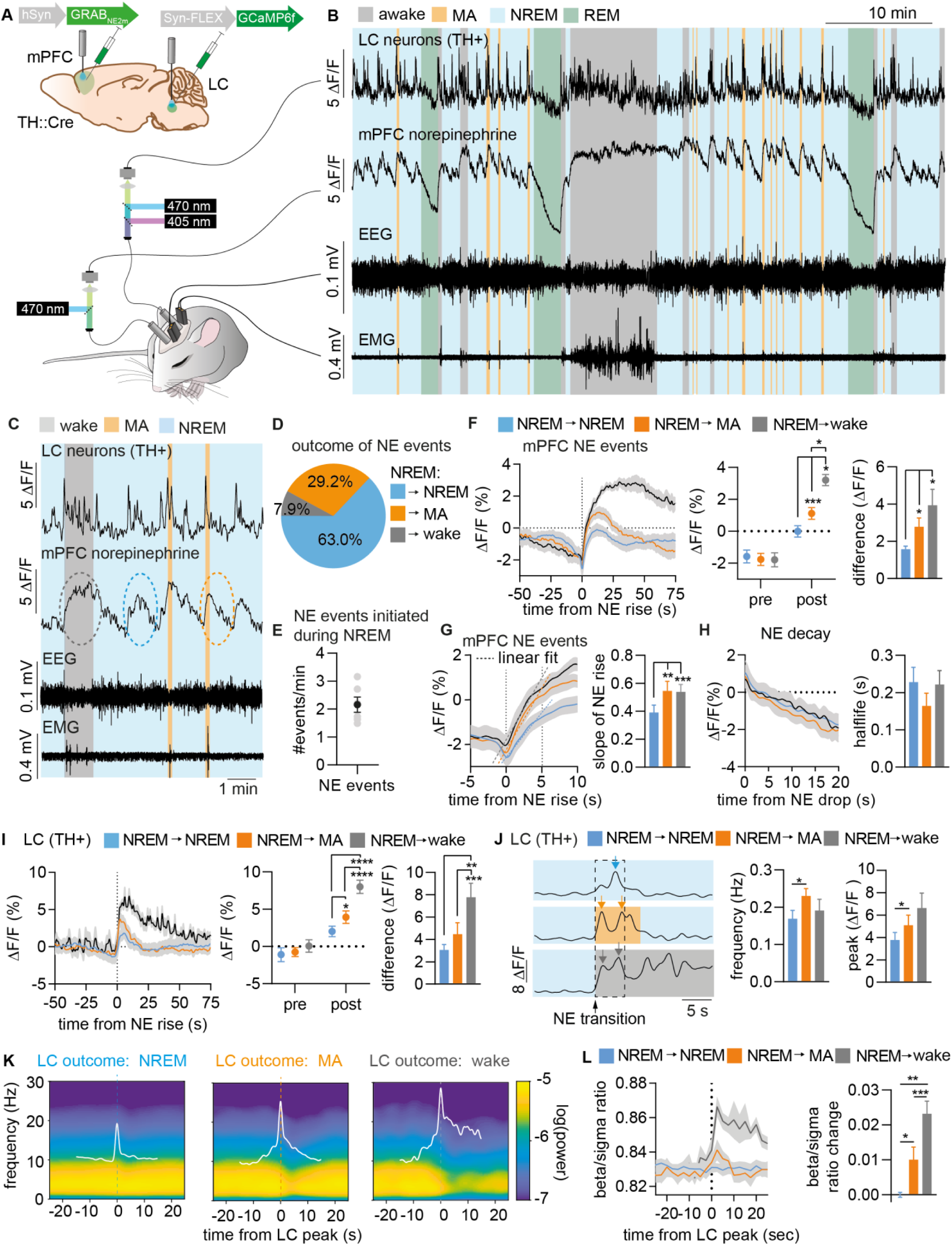
Prefrontal NE levels show dynamic fluctuation during NREM sleep. **A**. In TH::Cre mice, AAV5 viral vectors containing hSyn-GRAB_NE2m_ and syn-FLEX-GCAMP6f were injected into mPFC and LC, respectively, alongside implantation of optic fibers and screws for EEG and EMG recordings. **B**. Representative sample traces of simultaneous recordings of locus coeruleus (LC) activity, prefrontal norepinephrine (NE), EEG and EMG. Periods of wakefulness (grey), microarousals (MA) (orange), NREM (blue), and REM (green) are indicated. **C**. Representative traces showing NREM-related NE events associated with sustained NREM (blue circle), microarousal (MA) (orange circle), or waking (grey circle). **D**. Summary plots of outcome and (**E**) occurrence of NE events during NREM. **F**. Mean NE traces aligned to onset of NE rise (left) at behavioral transitions, mean NE levels prior to (−10 to 0 s, ‘pre’) and after (max value, 0-50 s, ‘post’) behavioral transition (middle), and difference between ‘post’ and ‘pre’ level of NE (right). **G**. The mean slope of initial NE increase (0-5 s). **H**. Mean NE decay leading up to transition (left) and summary of halflife (right). **I**. Mean LC traces aligned to onset of NE rise (left). ‘Pre’ (−10 to 0 s, mean value) and ‘post’ (0-50 s, max value) transition levels of LC (middle) and difference between LC activity (‘post’ minus ‘pre’) (right). **J**. Left: Representative LC traces with LC events (arrows) (0-5 s after NE transition) within LC activity bouts leading to sustained NREM, microarousal (MA) and wake. Mean frequency (middle) and accumulated peak values (right) depending on outcome. **K**. Mean spectrograms aligned to detected LC events during NREM, microarousal (MA) and wake. **L**. The mean ratio of beta and sigma power at onset of LC events (left) as well as the change in mean ratios across transition (before (−10 to 0 s) to after (0 to 5 s) (right). ANOVA, Paired t test and Wilcoxon test, n=6. Data is shown as mean±SEM. *p < 0.05, **p < 0.01, ***p < 0.001, ****p < 0.0001.

We expected to observe a constant level of prefrontal NE during NREM due to reported findings of tonic LC activity. Intriguingly, we observed a pronounced oscillatory pattern of prefrontal NE levels occurring every ∼30 s (Figure 1C and E, Suppl. Figure S4). Surprisingly, 63% of these NE events occurred during uninterrupted NREM sleep, whereas 29.2% and 7.9% resulted in microarousals and awakenings, respectively (Figure 1D). NE events associated with sustained NREM had a smaller trough-peak amplitude (Figure 1F) and slower initial increase (first 0-5 s) (Figure 1G) compared to NE events related to microarousal and waking. We saw no difference in the rate of NE decline leading up to different NREM transitions (Figure 1H). Prior to NE rise, we observed initiation of phasic LC activity bouts, whereas NE decline occurred following LC silence (Figure 1C); the amplitude of LC activity bouts correlated with the level of NE increase and arousal level (Figure 1I). The frequency and value of LC peaks (0-5 s) were reduced at sustained NREM compared to microarousals (Figure 1J). To confirm the behavioral state of the animal during the different NREM related LC/NE events, we mapped EEG activity at various state transitions. LC/NE events related to microarousal and wake transitions led to desynchronization and increased beta/sigma ratio, whereas LC/NE events associated with sustained NREM did not (Figure 1K-L), confirming that the majority of LC/NE events during NREM is not related to awakening.

Previously, single-electrode electrophysiological approaches demonstrated tonic LC activity during sleep; however, using a cell-specific and population-based imaging approach, we here show that phasic LC activity bouts are linked to a slow oscillatory pattern of prefrontal NE levels – the majority of which is an integral part of NREM sleep architecture.

### Low NE during NREM associates with spindle activity

Next, we sought to determine if the NE oscillations during NREM sleep reflect distinct phases of sleep related to sleep spindles: NREM stage 2 (N2) exhibits an increased occurrence of spindles (1-2 s of 8-15 Hz sigma activity in EEG), whereas NREM stage 3 (N3) display increased delta activity (0.5-4 Hz) (Andrillon et al., 2011). Considering previous reports of low LC activity during spindle activity (Aston-Jones and Bloom, 1981; Rajkowski et al., 1994; Swift et al., 2018), we investigated how NE levels change in the presence of sleep spindles. First, we plotted delta/sigma ratio, which computes contrasting values for N2 (low ratio) and N3 (high ratio) sleep stages. Accordingly, we observed an oscillatory pattern of delta/sigma ratio that inversely correlates with NE levels (Figure 2A); NE peaks occurred when delta/sigma ratio was at its maximum (Figure 2B), while NE troughs correlated with the lowest point of delta/sigma ratio (Figure 2C). Next, we used the EEG to detect spindle occurrences during sleep (Figure 2D-E) to evaluate NE levels at spindle onset. Spindle onset correlated with NE troughs and with reduced delta/sigma ratio (Figure 2F). Detection of high delta epochs during NREM sleep did not correlate with NE levels (Suppl. Figure S5). These findings suggest that the oscillatory pattern of NE during NREM underlies different sleep periods with low NE levels possibly having a permissive effect on spindle occurrences, while higher NE levels are associated with low spindle occurrences and lower amount of sigma activity. Next, we mapped prefrontal neuronal Ca^2+^ activity at NE troughs during NREM by co-expressing jRGECO1A and GRAB_NE2m_ in mPFC neurons (Figure 2G). Neuronal Ca^2+^ activity peaked when NE was at its lowest, and drastically declined when NE started to rise (Figure 2H). Thus, high neuronal Ca^2+^ activity coincides with periods of spindle activity and low delta/sigma ratio (Figure 2I). These results indicate that neuronal Ca^2+^ level reflects periods of high sigma power (Seibt et al., 2017).

**Figure 2.**
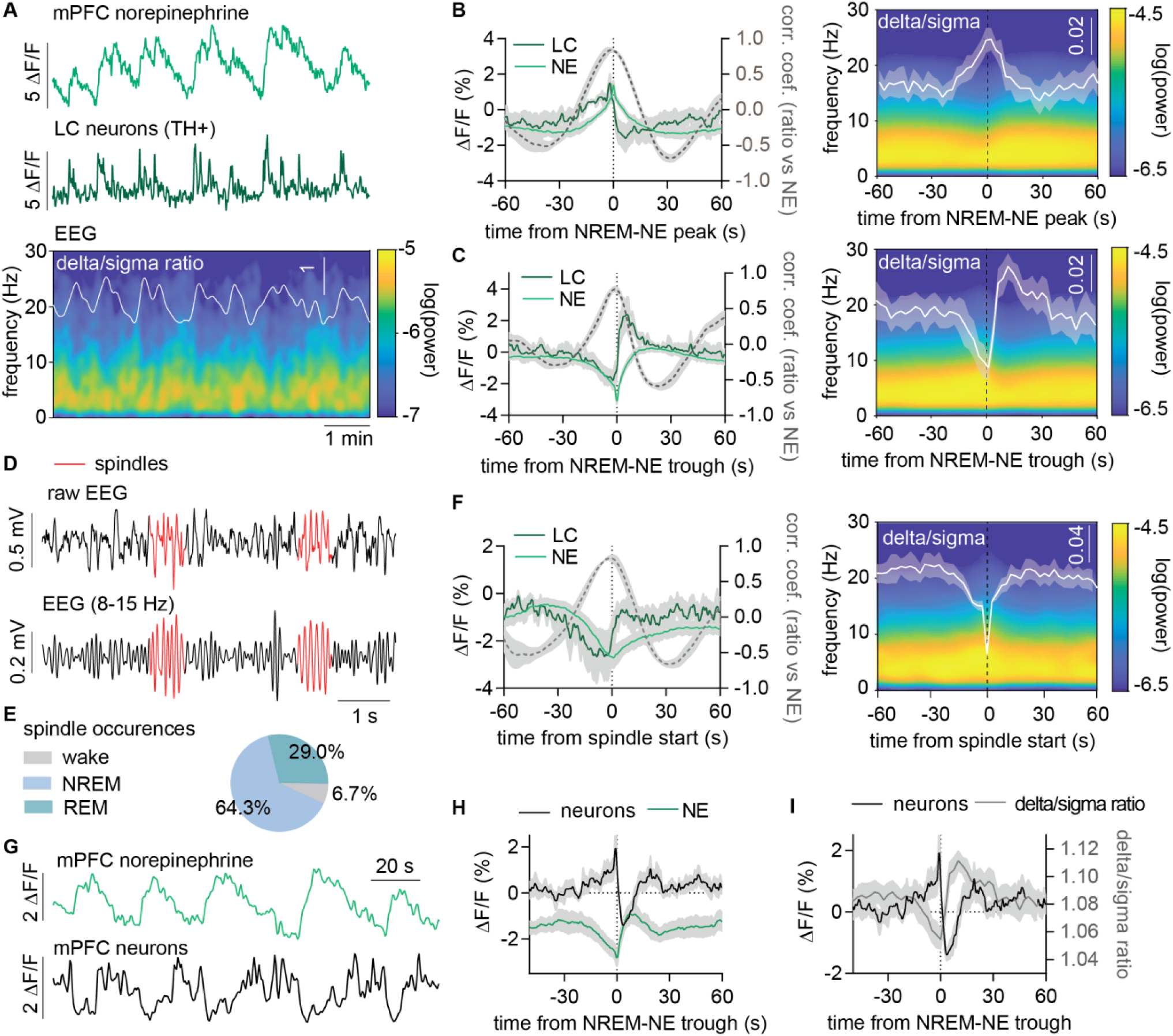
Low NE during NREM associates with spindle activity. **A**. Representative traces from simultaneous measurements of NE and LC during NREM and their association with EEG spectrogram and delta/sigma ratio. **B+C**. Left: Mean LC (dark green) and NE (green) traces aligned to time of NE peak (B) or trough (C) during NREM. The dotted grey trace is the correlation coefficient between delta/sigma ratio (shown in right part of figure) and NE level. Right: The mean spectrogram and delta/sigma ratio aligned around NREM-NE peak (B) or trough (C). **D**. Example trace showing the detection of spindles (red) from the EEG trace. **E**. The percentages of spindles occurring during wake, NREM and REM. **F**. Mean LC, NE traces and correlation coefficient (delta/sigma ratio versus NE levels) at spindle onset during NREM (left). The right panel shows the mean spectrogram and delta/sigma ratio around spindle onset. **G**. Representative traces of simultaneous measurements of prefrontal NE and neuronal Ca^2+^ during NREM. **H**. Mean neuronal and NE traces around NE minimum at sustained NREM sleep. **I**. Mean neuronal trace and delta/sigma ratio around NE minimum. n = 5. Data is shown as mean±SEM.

### REM is characterized by prolonged continuous NE drop

Next, we investigated NE dynamics during REM. Interestingly, we observed the largest dynamic range of NE during REM (Figure 3A and B). At the transition from NREM to REM, LC phasic events cease and NE levels drop dramatically (Figure 3C). Likewise, at the offset of REM sleep, LC shows phasic activity accompanied by a huge increase in NE (Figure 3C) that was similar between different behavioral outcomes (microarousal or wakefulness) (Suppl. Figure S7). The magnitude of the NE drop during REM sleep correlated with the REM duration (Figure 3D). Further, we discovered that changes in LC activity and NE precede both REM onset and offset (Figure 3E-F). The level, drop magnitude, and duration of the pre-REM drop in NE did not correlate with the subsequent duration of REM (Suppl. Figure S6). We determined the mean delay from when NE started dropping until REM onset (42.3 ± 3.8 s), as well as the delay from NE increase to REM offset (4.7 ± 0.5 s) (Figure 3G). We then aligned LC and NE traces to the time point at which NE starts declining (Figure 3H), and saw that EEG power in the sigma range increased within the first ∼40 s of NE decline leading up to REM onset, while delta frequency power decreased (Figure 3I-J) illustrated by a pronounced reduction in delta/sigma ratio (Figure 3K). Following the onset of REM, no change in delta/sigma ratio is observed, but instead theta/sigma ratio builds up (Figure 3L), which is also reflected by increased neuronal activity (Figure 3M, Suppl. Figure S7). Finally, we speculated if the initial NE drop differed between NREM and REM, and found it to decrease slightly faster when transitioning to REM sleep (Figure 3N and O).

**Figure 3.**
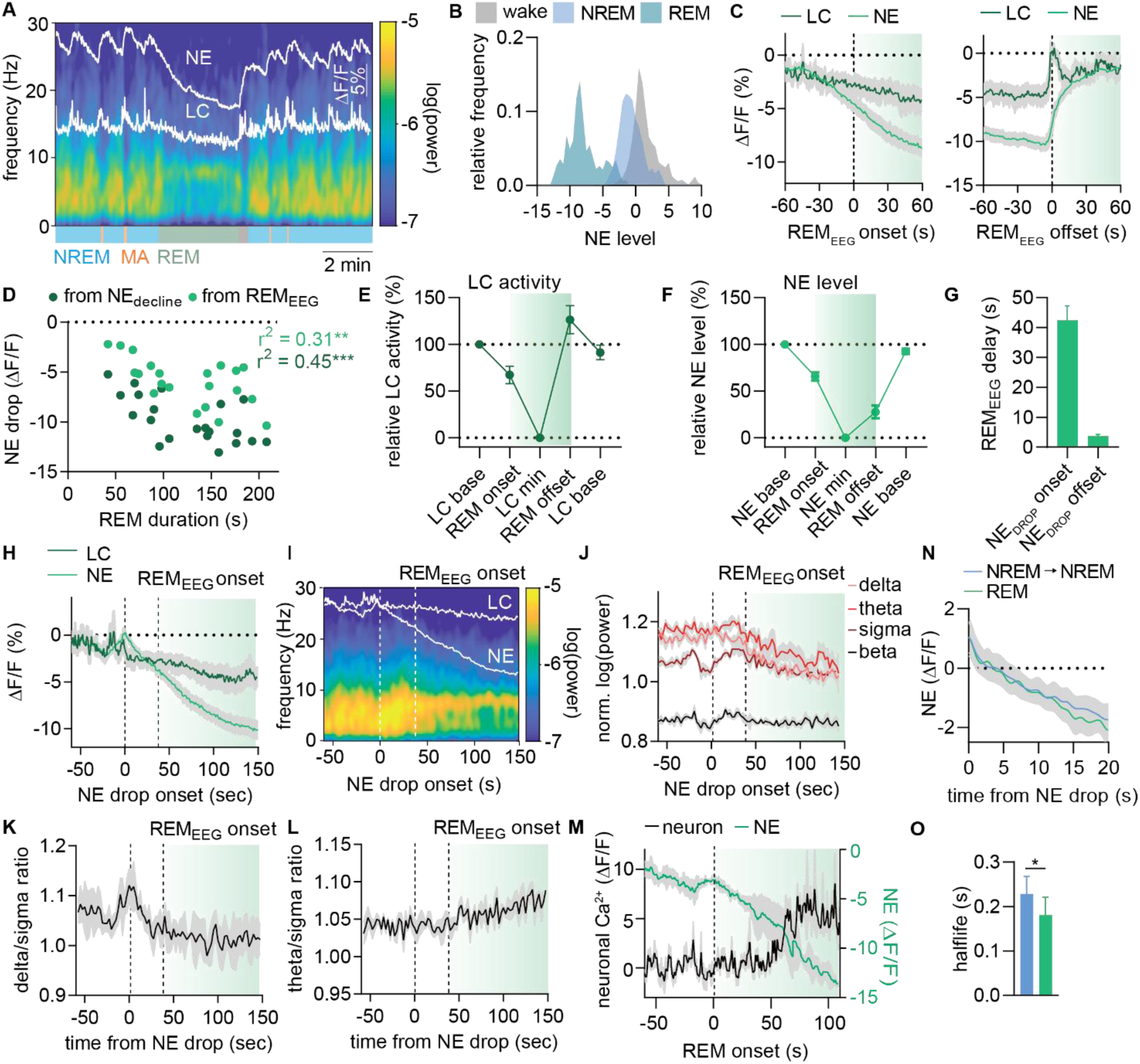
Drop in LC-NE activity precedes transition to REM sleep and is associated with increased delta/sigma ratio. **A**. Representative LC and NE traces surrounding a REM bout with associated EEG power spectrum. **B**. Mean NE dynamics during wake, NREM and REM. **C**. Mean LC and NE traces aligned to onset and offset of REM sleep. **D**. Correlation between REM bout duration and NE drop measured from the start of NE decline (dark green) or from start of REM (light green). **E+F**. Relative LC activity (E) and NE level (F) at baseline, REM onset, minimum level, REM offset, and post-REM baseline. **G**. Mean delay from onset of NE drop to REM onset and from NE rise to REM offset. **H-J**. Mean LC activity and NE level (H), EEG power spectrum (I), and filtered band power traces (J) aligned to onset of NE drop. **K+L**. Mean ratios of delta/sigma (K) and theta/sigma (L) aligned to onset of REM drop. **M**. Mean neuronal Ca^2+^ activity and extracellular NE level in mPFC aligned to REM onset. **N**. Mean of initial 20 s of NE drop leading to REM sleep and continued NREM sleep. **O**. Decay time of NE drops resulting in continued NREM or REM. Wilcoxon test, n=6. Data is shown as mean±SEM. *p < 0.05, **p < 0.01, ***p < 0.001.

### Astrocytes are silent during sleep despite LC-NE activity and is recruited upon awakening

Astrocytic Ca^2+^ is tightly linked to LC-NE activity via α1 activation, and astrocytes have recently been described to have diminished Ca^2+^ activity during sleep (Bojarskaite et al., 2020). On the other hand, our present observations demonstrate LC-NE activity during slow wave sleep. We therefore set out to study the association between astrocytic Ca^2+^ and LC-NE during the sleep-wake cycle. We performed dual-color imaging of astrocytic Ca^2+^ and extracellular NE in mPFC with simultaneous recording of LC neuronal Ca^2+^ activity during natural sleep (Figure 4A-C). We confirmed astrocytes to be silent during sleep (Figure 4D). Remarkably, when aligning astrocytic traces to detected LC events during sleep, we observed no astrocytic response though NE increased (Figure 4E and G), while LC events during wakefulness elicited both NE and astrocytic events. (Figure 4F and G) also evident from the observation that the positive cross correlation between LC and astrocytic Ca^2+^ during wakefulness is completely absent during NREM sleep (Figure 4H).

**Figure 4.**
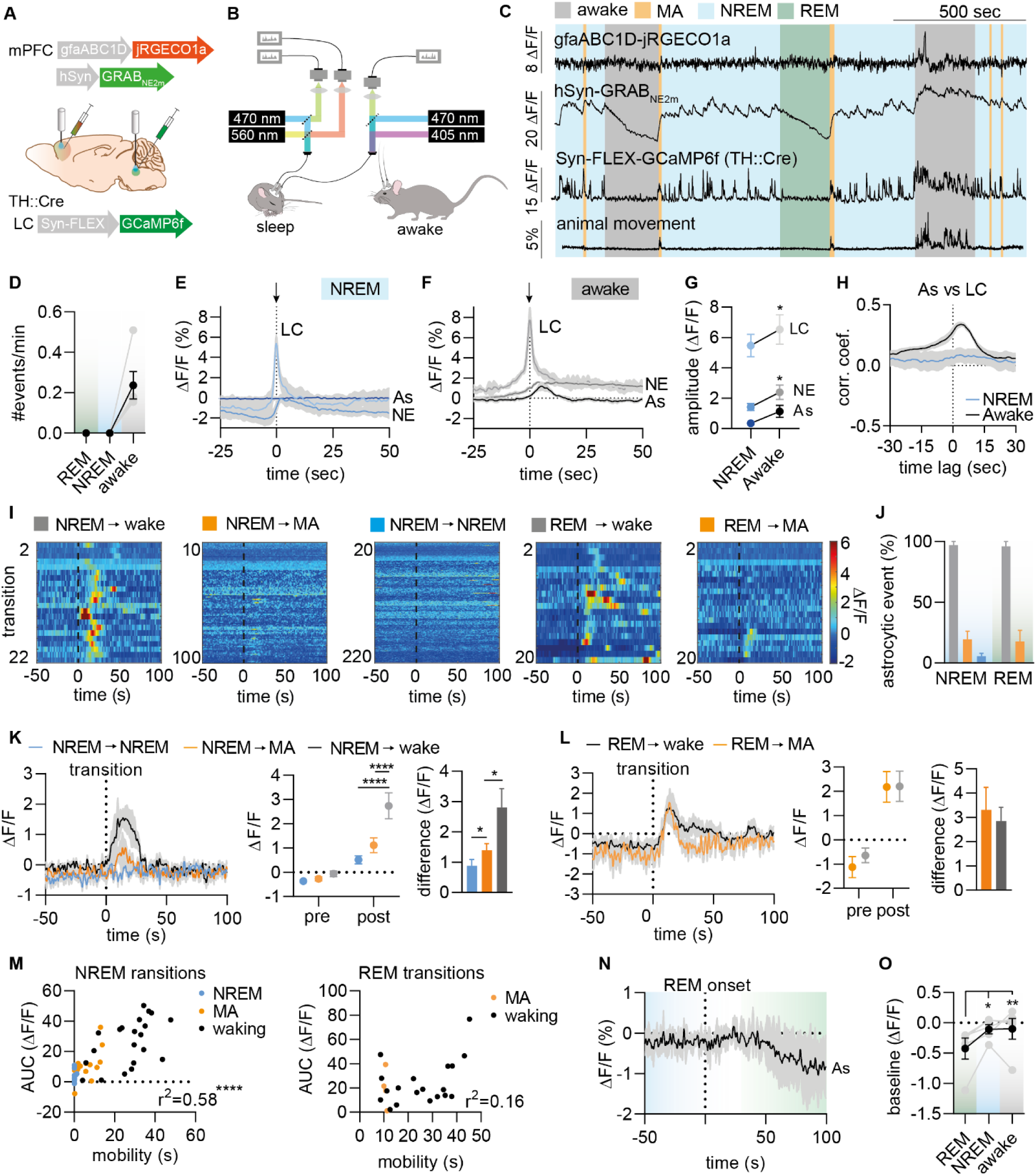
Astrocytes are silent during sleep despite LC-NE activity and is recruited upon awakening. **A+B**. The norepinephrine biosensor, GRAB-NE_2m_, and red-shifted calcium indicator, jRGECO1a, under a GFAP promoter was co-injected in mPFC, while GCaMP6f was expressed in TH positive neurons within LC. Fiber implants were inserted above mPFC and LC for fiber photometric recordings. **C**. Example traces showing NE, astrocytic Ca^2+^ and LC Ca^2+^ activity across sleep-wake cycles. **D**. No astrocytic events were detected during NREM and REM. **E+F**. Mean LC events selected during NREM (E) and wakefulness (F) with the corresponding changes in NE and astrocytic activity. **G**. Summary plot of LC events and corresponding NE and astrocytic activity. **H**. The correlation coefficient between LC and astrocytic activity during wakefulness and NREM states. **I**. Heatplots showing astrocytic Ca^2+^ activity aligned to transitions from NREM and REM **J**. Percentage of events leading to astrocytic response. **K**. Left: mean astrocytic response aligned to initiation of NE rise leading to either NREM, microarousal (MA) or awakenings. Middle: Astrocytic baseline levels (pre) and peak values (post) across transitions. Right: Difference between post and pre values. **L**. Mean astrocytic response and summary plots during transition from REM to either wakefulness or MA. **M**. Correlation between mobility post-transitions and area under the curve (AUC) of astrocytic Ca^2+^ responses for NREM (left) and REM (right) transitions. **N**. Mean astrocytic Ca^2+^ aligned to onset of REM. **O**. Astrocytic Ca^2+^ baseline during REM, NREM, and wakefulness. ANOVA, Paired t test, Wilcoxon test, n=5. Data is shown as mean±SEM. *p < 0.05, ****p < 0.0001.

Next, we looked at astrocytic activation upon transitions from NREM and REM (Figure 4I); astrocytes responded consistently upon awakening, upon 20% of microarousals and at almost none of sustained NREM transitions (Figure 4J). When responding, astrocytic events had a higher amplitude upon awakening from NREM compared to a microarousal (Figure 4K), but the amplitude was similar between REM transitioning to awakening or microarousal, (Figure 4L). The magnitude of astrocytic responses to transitions from NREM correlates with the amount of mobility that follows the transition (Figure 4M), indicating that the astrocytic response to NE is not all-or-nothing, but depend on the level of NE. Interestingly, this correlation is absent at REM transitions (Figure 4M) possible due to the big dynamic range of the NE increase. Lastly, following the onset of REM sleep, astrocytic Ca^2+^ baseline activity decreases slightly, which could be related to REM-mediated NE decline (Figure 1N-O).

### Astrocytes are less responsive to NE increases during sleep and anesthesia compared to wake

The absence of astrocytic Ca^2+^ response to NE increases during NREM was unexpected. To explore this, we conducted 10 s (5 Hz 10 ms pulses) of blue light stimulation of channelrhodopsin2 (ChR2)-expressing LC neurons during sleep and wakefulness, while simultaneously measuring prefrontal astrocytic Ca^2+^ (Figure 5A). Light intensity was adjusted to elicit an astrocytic Ca^2+^ response during the awake condition. When stimulated during wakefulness, ChR2-expressing mice would arrest their ongoing behavior, while stimulation during sleep would eventually awaken the animals (Figure 5B). Interestingly, the astrocytic Ca^2+^ response to LC stimulation during sleep had a slower increase compared to the wake condition (Figure 5D-G). Astrocytes responded with similar amplitude and kinetics to both wake and sleep stimulations, but the baseline undershoot observed after the end of stimulation in the wake condition was not observed during sleep (Figure 5D-G). We did not observe any astrocytic activation or behavioral effect from light stimulation in the eYFP injected control group (Figure 5C and 5E). Our results indicate that astrocytes display a slower recruitment of Ca^2+^ in response to NE elevations during sleep versus awake conditions. However, we cannot rule out that the difference in astrocytic response is due to different pre-stimulation NE levels in sleep versus awake conditions. To exclude this possibility, we conducted an experiment that allowed us to circumvent the amount of endogenous NE levels by pharmacological means. Using GLT1-GCaMP7 mice (Monai et al., 2016), we imaged somatosensory cortical astrocytic Ca^2+^ under a 2-photon microscope while locally puffing on the α1 noradrenergic receptor agonist, methoxamine (50 μM) (Figure 5H). We delivered methoxamine during wakefulness and then administered either ketamine/xylazine (KX) or saline systemically and repeated the methoxamine puff (Figure 5I and J). KX reduced the propagation area of the astrocytic Ca^2+^ responses compared to wake; an effect not observed in the saline control group (Figure 5K). Together, these results indicate that the responsiveness of astrocytic Ca^2+^ response to NE-mediated activation is reduced during sleep compared to wake.

**Figure 5.**
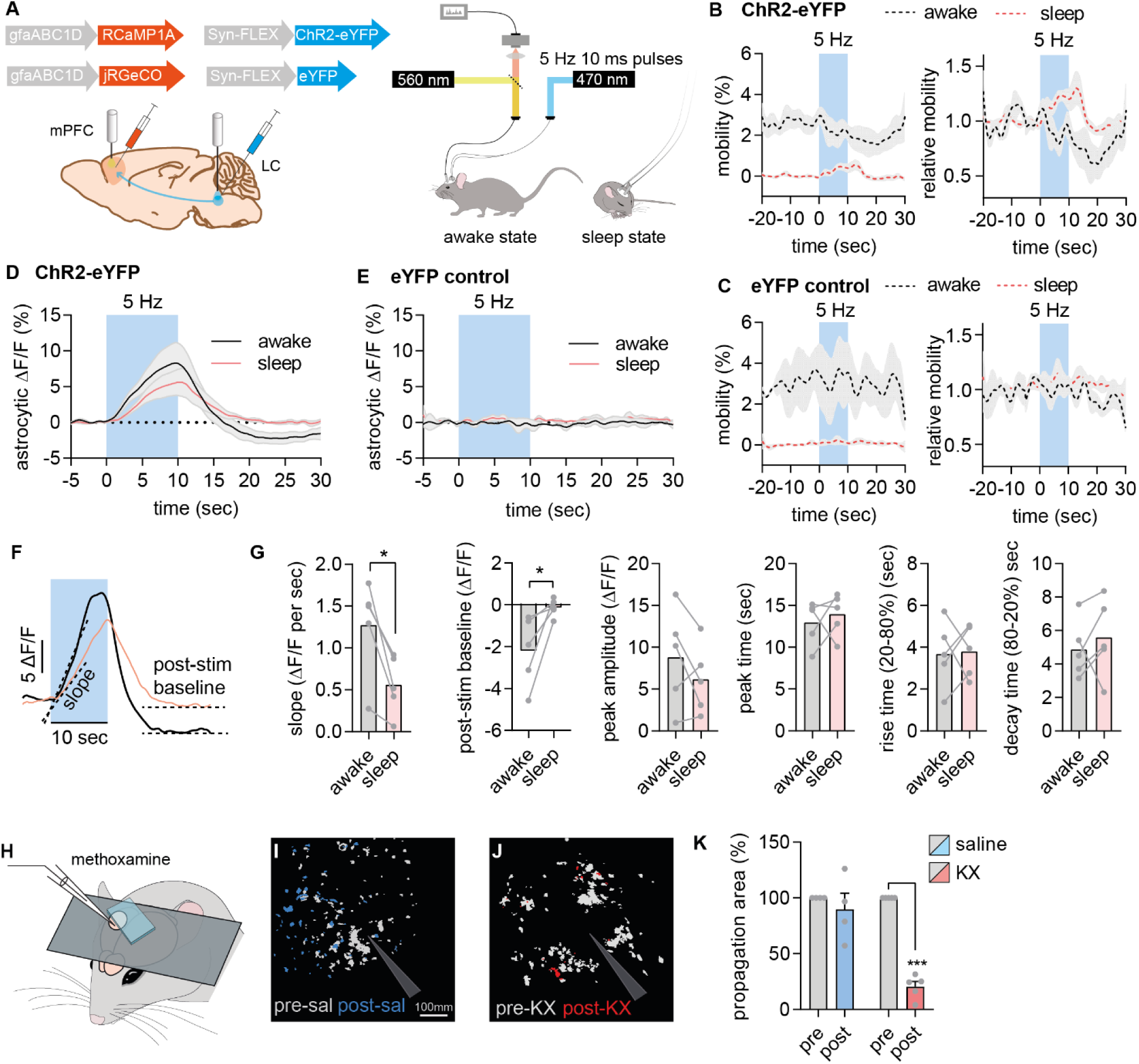
Astrocytes are less responsive to NE during sleep and anesthesia compared to wake. **A**. We expressed RCaMP1A or jRGECO1a in prefrontal astrocytes in mPFC and ChR2-eYFP or eYFP under the Syn-FLEX promoter bilaterally in LC of TH-Cre animal in order to stimulate LC, while measuring prefrontal astrocytic Ca^2+^ during sleep/wake patterns. **B-C**. Mobility and mobility relative to baseline prior to and after 10 s of 5 Hz optogenetic stimulation (light blue) during sleep and wakefulness in ChR2-eYFP injected animals and eYFP injected controls. **D-E**. Astrocytic Ca^2+^ in response to optogenetic stimulation during sleep and wakefulness in ChR2-eYFP injected animals and eYFP controls. **F-G**. Kinetics of the mean astrocytic Ca^2+^ response to optogenetic stimulation during sleep and wakefulness in ChR2-eYFP animals. **H**. Schematic diagram of the experimental setup in awake head-restrained mice. **I-J**. Representative examples of increases in GCaMP7 signal 0-60 s after local administration of methoxamine. Light gray indicates the pixels within the field of view that increased in response to methoxamine, whereas the red and blue map the pixels that increased 15 min after administration of either KX or saline i.p within the same field of view. **K**. Summary plots comparing the relative area of GCaMP7 increases following methoxamine injection before and after administration of KX. Optogenetic experiment: n=5. Puff experiment: saline, n = 4; KX, n = 5. Data is shown as mean±SEM. Paired t test and Wilcoxon test. *p < 0.05 **** P < 0.0001.

## Discussion

The dynamic relationship between LC activity and cortical NE levels during sleep is unknown and limits our understanding of how NE shapes sleep processes. We show for the first time that periodic LC activity bursts during NREM sleep lead to a slow oscillatory pattern of prefrontal NE levels (∼every 30 sec) with the majority of NE increases happening during sustained sleep, while a smaller proportion result in microarousals and awakenings. NE troughs during NREM sleep coincide with spindle occurrences that are immediately terminated at initiation of NE rise. Prolonged continuous NE decline (>40 s) sets the stage for REM onset and build-up of theta oscillations, while REM offset is preceded by a steep increase in NE. The NE increases during NREM sleep do not lead to astrocytic Ca^2+^ responses; rather, the positive correlation between LC and astrocyte activity during wakefulness is non-existing during NREM with less astrocytic responsiveness to NE during sleep and anesthesia compared to wakefulness. These findings suggest that the role of wake-promoting systems are not limited to awake behavior, but also play a regulatory role in integral sleep processes.

### Phasic versus tonic LC activity during NREM sleep

We show that NREM sleep is characterized by periodic (∼every 30 sec) bouts of LC Ca^2+^ increases that is likely caused by LC phasic activity bursts of which the majority did not lead to microarousals or awakenings. Rat and monkey studies have reported a slow tonic activity pattern of LC during NREM sleep (Aston-Jones and Bloom, 1981; Eschenko and Sara, 2008; Eschenko et al., 2012; Foote et al., 1980; Rasmussen et al., 1986), but this was not evident from our recordings. It is possible that single or a few LC spikes will not increase internal Ca^2+^ levels to an extent where we can pick it up in our LC measurements. In addition, tonic LC activity during NREM is not reflected in our prefrontal NE signal; rather, NE declines between LC activity bouts. In anesthetized rats, electrical stimulation of LC increases cortical NE levels in a linear frequency-dependent manner and correlates with desynchronization of EEG (Berridge and Abercrombie, 1999; Berridge et al., 1993; Florin-Lechner et al., 1996). Furthermore, these studies show that a similar number of LC spikes release NE more efficiently by a bursting compared to tonic discharge, which may explain why the reported tonic spiking pattern of LC neurons are not evident from our NE measurements. However, species-dependent differences in NREM-related LC firing might also explain these discrepancies; in a pioneering mouse study performing single unit recordings in LC, LC neurons are silent during both NREM and REM sleep and only active during awakenings or awake behavior (Takahashi et al., 2010).

None of the single unit studies report LC burst firing during NREM, though it is possible that our observed phasic NREM LC events are the result of recruitment of a larger proportion of available LC neurons. It is conceivable that the Ca^2+^ imaging approach of fiber photometry detects population-based events more reliably than single unit recordings.

### NE increases during NREM

We observe that the majority of NE increases lead to sustained sleep (∼63%), whereas a smaller proportion leads to microarousals (∼29%) and awakening (∼8%). The degree of LC activation and NE increase are reflected in the behavioral outcome with larger LC event amplitude and steeper NE increase associated with microarousal and awakening compared to sustained sleep. The NE slow oscillations during NREM created by periodic activity bouts of LC neurons might be a survival mechanism creating windows of increased sensitivity towards sensory inputs and abrupt awakenings in case of threats. This is supported by findings in rats showing that higher levels of tonic LC activity during NREM increase the likelihood of sound-evoked awakenings (Hayat et al., 2020). Also, mice with NE deficiency display reduced sleep latency and reduced sensory-evoked awakenings (Hunsley and Palmiter, 2004).

### Does NE underlie NREM sleep stages?

We show that cortical spindles are associated with NE troughs and vice versa. This fits well with previous reports of LC silence prior to spindles (Aston-Jones and Bloom, 1981; Rajkowski et al., 1994; Swift et al., 2018) and LC activity happening at spindle termination (Swift et al., 2018). Our findings suggest that NE needs to go below a certain threshold in order to permit spindles. Spindles are generated by thalamocortical oscillations generated by the interplay between thalamic reticular neurons and the thalamocortical projecting neurons. It has been suggested that reductions in NE allow the rhythmic up and down states of the thalamocortical neurons, while increases in NE generate a constant upstate abolishing the spindle oscillations (Lee and McCormick, 1996). Our findings are in support of these electrophysiological studies. It is possible that NE fluctuations during NREM sleep could underlie transition between sleep stages with low NE levels corresponding to spindle-rich NREM stage 2 sleep. A study suggests that single LC spikes correlate with the initiation of cortical slow-wave up-states during NREM (Eschenko et al., 2012) suggesting that NE levels might be necessary for upregulation of delta generation during NREM stage 3 (slow-wave) sleep. However, we did not observe any correlations between epochs of high delta activity and NE levels.

### NE declines continuously during REM

While LC silencing during REM sleep is established, cortical NE levels during REM has remained unaddressed. We demonstrate for the first time that there is a monotonic NE decrease throughout the REM episode. Furthermore, prior to REM onset (∼40 s), NE starts to decline. This period is characterized by increased sigma activity indicative of spindle activity. The occurrence of spindles just prior to REM is well-known in sleep literature and are categorized as intermediate sleep or NREM stage 2 sleep (Andrillon et al., 2011). We show that this phase is characterized by NE decline similar to the periodic NE troughs during NREM, which are also characterized by spindles. However, at some point, the decline of NE goes below a certain threshold leading to an abolishment of spindles and a buildup of theta activity. Thus, NE decline is not always linked to spindle activity, but seems permissive for REM induction. This is supported by studies showing that mild electrical stimulation of LC or pharmacological enhancement of NE transmission reduces REM episodes (Gervasoni et al., 2002; Jones, 1991; Singh and Mallick, 1996), while selective neurotoxic lesioning of LC-NE neurons increases REM episodes (Monti et al., 1988). However, neither optogenetic inhibition of LC nor CRISPR knockdown of NE-producing enzyme within LC altered REM sleep (Carter et al., 2010; Yamaguchi et al., 2018) indicating that other REM-inducing components are also needed.

### Prefrontal neuronal Ca^2+^ increases occur at spindle and theta activity

We observed a pronounced upregulation of prefrontal neuronal Ca^2+^ during NE troughs in NREM sleep as well as during REM sleep. During transitioning into REM sleep, cortical neurons have been demonstrated to go from a slow delta oscillatory pattern to a consistent up-state characterized by depolarization (Steriade et al., 2001). Recent studies show that spindle activity and REM sleep are associated with increased Ca^2+^ entry specifically at the dendrites of cortical neurons, not the soma (Li et al., 2017; Seibt et al., 2017). Furthermore, the Ca^2+^ activity did not correlate with delta oscillations (Seibt et al., 2017). Thus, the neuronal Ca^2+^ upregulations we observe are probably due to Ca^2+^ entry at the dendrites and associates with oscillations in the sigma and theta frequency ranges. Since these rhythms are tightly linked to memory consolidation possibly through synaptic remodeling (Ulrich, 2016), the neuronal Ca^2+^ increases happening at low NE levels might be involved in these synaptic processes. The fact that the neuronal activity is abruptly ended when NE increases, supports the permissive effect of NE on memory consolidation.

### The involvement of astrocytes in sleep

Our results indicate that prefrontal astrocytes are part of the wake-promoting system with lack of astrocytic Ca^2+^ responses during sleep and consistent Ca^2+^ increases upon awakenings that positively correlates with the degree of awakening. Similar findings have been made in recent studies conducting two-photon or head-mounted epifluorescence microscope imaging of astrocytic Ca^2+^ responses in sleeping mice (Bojarskaite et al., 2020; Ingiosi et al., 2019). Our result show that astrocytic Ca^2+^ levels do not reflect the dynamic NE fluctuations during NREM sleep; however, we observed a decline in astrocytic Ca^2+^ baseline during REM sleep, where NE declines drastically. The lack of astrocytic response to LC activity bouts was surprising, since cortical astrocytes have been shown to elevate Ca^2+^ predominantly in the presence of bursting NE axons (Oe et al., 2020). By stimulating astrocytes with either optogenetically induced elevations of endogenous NE or pharmacological activation of α1 noradrenergic receptors and comparing the response between wakefulness and sleep or anesthesia, we observed a reduced responsiveness of astrocytes towards NE. These results indicate that astrocytes might play a role in sleep maintenance by decoupling from the NE increases. Several studies have reported the effect on sleep of compromising IP_3_-mediated Ca^2+^ signaling constitutively in astrocytes although the involvement of IP_3_ signaling in sleep remains controversial (Cao et al., 2013). One study shows increased duration of REM sleep and related hippocampal theta oscillations (Foley et al., 2017), while another study reported no effect on REM sleep (Bojarskaite et al., 2020). Furthermore, in anaesthetized animals, impairment of astrocyte-related exocytosis reduced the theta synchronization between hippocampus and mPFC (Sardinha et al., 2017). Bojarskaite et al., 2020 also reports more frequent and shorter NREM related sleep stages with a dysfunctional increase in spindle occurrences. These results suggest that astrocytes are required for intact sleep, but further investigations are needed in order to clarify this.

## Conclusion

In summary, our results provide evidence that LC-NE and astrocytic fluctuations not only promote shifts in brain states during wake adaptive behavior, but also shape critical sleep processes. This illustrates the versatility of wake-promoting systems, where both up- and downregulations play an active role in sleep state transitions. Future studies may further advance our understanding of the pathophysiology of neuropsychiatric disorders characterized by dysfunctional LC-NE signaling and sleep disturbances and sleep-targeted interventions may represent new avenues for treatment.

## Materials and Methods

### Mice

Wildtype C57BL/6 mice were acquired from Janvier Labs at 7 weeks of age. Heterozygous TH::Cre mice were bred on a C57BL/6 background. Glt1-GCaMP7 mice were obtained from RIKEN BioResource Research Center, Japan, and bred in-house on a C57BL/6 background. Glt-1-GCaMP7 mice express GCaMP7 in 95.2% of cortical astrocytes with minimal expression in other glial cell types and about 50% cortical neurons in L4 and L6. All mice used in this study were male. Animals were housed with unlimited access to food and water in a normal 12-h light/dark cycle. Animals were 16-20 weeks old at time of behavioral assessment. All experiments conducted at University of Copenhagen were approved by the Danish Animal Experiments Inspectorate and were overseen by the University of Copenhagen Institutional Animal Care and Use Committee (IACUC), in compliance with the European Communities Council Directive of 22 September 2010 (2010/63/EU) legislation governing the protection of animals used for scientific purposes. All animal experiments conducted at University of Rochester was approved by the University Committee on Animal Resources of the University of Rochester.

### Surgery

All surgeries were performed in accordance with institutional guidelines. Mice were 7-15 weeks old at time of surgery. General anesthesia was induced using 5% isoflurane, and afterwards maintained at 1-3% isoflurane. The mice were placed in a stereotactic frame and received preoperative buprenorphine (0.05 mg/kg) for general analgesia along with lidocaine (0.03 mg/kg) at the incision site. An incision was made on the scalp between the ears and the skull was aligned. Four burr holes were drilled in the skull with an electrical drill (Tech2000, RAM Microtorque 45,000 rpm) according to stereotactic coordinates relative to bregma. In mPFC, we co-injected AAV9 encoding GRAB_NE2m_ under the neuronal hSyn promoter (provided by Yulong Li) and AAV/PHP.eB encoding an enhanced GFAP promoter, GfaABC1D, driving jRGECO1a. Simultaneously, we injected AAV5 encoding floxed GCaMP6f in LC in a TH-Cre driver mouse line. In a separate group of mice, AAV5 encoding jRGECO1a under a hSyn promoter was co-injected with the NE biosensor. Stereotactic coordinates used for mPFC injections: A/P +1.7 mm, M/L −0.3 mm, and D/V −2.00, −2.25, −2.50, and −2.75 mm (125 nL virus infused at each depth) and for LC injections: A/P −5.5 mm, M/L: −0.9 mm, and D/V: - 3.2 mm, −3.4 mm, - 3.6 mm, and −3.8 mm (125 nL at each depth). Virus was infused at 100 nL/min and the needle was left in place for additional 7 minutes and then slowly withdrawn. 0.8 mm low impedance stainless steel screws (NeuroTek) were screwed into two burr holes located above frontal cortex (contralateral side from the optic implant), and above the cerebellum (reference area). A silver wire (W3 Wires International) was inserted into the trapezius muscle serving as an electromyogram (EMG) electrode. Mono fiber-optic cannula (400 μm,0.48 NA, Doric Lenses) attached to a 2.5-mm diameter metal ferrule was then implanted in mPFC (A/P 1.7 mm, M/L −0.3 mm, D/V −2.50) and LC (A/P −5.5 mm, M/L: −0.9 mm, D/V: −3.65 mm) and cannulas and screws were fixed to the skull using dental cement (SuperBond). Prior to waking up, animals received carprofen (5 mg/kg) s.c.. A minimum of two weeks were allowed for sufficient expression and recovery.

For optogenetic LC experiments, AAV5 encoding floxed ChR2-eYFP or eYFP were injected bilaterally in LC at A/P −5.5 mm, M/L: +/-0.9 mm, D/V: −3.75 mm (300 nL virus) and dual fiber-optic cannula (200 μm, 0.22 NA, Doric Lenses) were implanted above LC (D/V: −3.65 mm).

| Construct | Company/provider | Cat# |
| --- | --- | --- |
| pAAV9-hSyn-GRAB<sub>NE2m</sub> | Yulong Li lab |  |
| AAV5-Syn-FLEX-GCaMP6f-WPRE-SV40 | Addgene | 100833-AAV5 |
| AAV/PHP.eB-GFAP-NES-jRGECO1a | Vector Biolabs | Custom made |
| AAV9-hSyn-jRGECO1a-WPRE | Addgene | 100854-AAV1 |
| AAV5-EF1a-DIO-hChR2(H134R)-EYFP | UNC Vector Core |  |
| AAV5-EF1a-DIO-EYFP | UNC Vector Core |  |

Mice used for *in vivo* two-photon imaging were 8 weeks old Glt-1-GCaMP7 mice. Mice were anesthetized under 2% isofluorane. Briefly, mice had head-plate implantation and a 2 mm craniotomy (with the dura carefully removed) was opened over the somatosensory cortex (AP: −0.8 mm, ML: +3.5mm). Brain tissue was carefully covered with a glass coverslip with 1/4 of the window space left open for insertion of a microelectrode. The cranial window was covered with aCSF during the surgical and imaging processes. Imaging was not started until more than 1hr after withdrawal of isoflurane. Intraperitoneal catheterization was prepared for injection of ketamine/xylazine (KX) (100/10 mg/kg, i.p.) and saline in the same volume as KX.

### Fiber photometry

Two pairs of excitation LEDs (560 nm and 465 nm; 465-nm and 405-nm, Doric Lenses, Tucker Davis Technologies) were connected to each their minicube (Doric Lenses) by attenuator patch cords (400-μm core, NA = 0.48, Doric Lenses). The minicube optics allow for the simultaneous monitoring of two spectrally separated fluorophores, with dichroic mirrors and cleanup filters chosen to match the excitation and emission spectra. LEDs were controlled by LED drivers (Thorlabs, Doric, Tucker Davis Technologies) and connected to a RX-8 or RX-10 real-time processor (Tucker-Davis Technologies). In mPFC, 560 nm and 465 nm excitation light were delivered through the same patch cord to stimulate jRGECO1a and GRAB_NE2m_ fluorescence, respectively. In LC, using the same patch cord, excitation light at 465 nm stimulated GCaMP6f fluorescence and 405 nm was an excitation isosbestic wavelength for GCaMP6f correcting for bleaching and signal fluctuations due to movement. 560nm/465nm and 465nm/405nm excitation were both sinusoidally modulated at 531Hz/211 Hz. Fiber-optic patch cords (400-μm core, NA = 0.48, Doric Lenses) provided a light path between the minicubes and the animals. Zirconia sleeves were used to attach fiber-optic patch cords to fiber implants on the animal.

Each of the four modulated signals generated by the four LEDs were independently recovered using standard synchronous demodulation techniques implemented on the RX-8/RX-10 real-time processor (sampling rate of 1000 Hz). The commercial software Synapse (Tucker-Davis Technologies) was used to control the signal processor and were aligned to video and EEG/EMG signals through in- or outcoming TTL pulses. Files were exported for analysis to MATLAB (MathWorks). ΔF/F calculations were based on the fitted 405 nm signal or by using the median of the fluorescence signal itself.

### Optogenetic stimulation

A 465 nm laser (SLOC Lasers) was used to generate 10 s stimulations (5 Hz 10ms pulses, 10 mW at the tip of the fiber) bilaterally in LC during states of quiet wakefulness and sleep simultaneous with mPFC fiber photometry recordings. Mice were left undisturbed in their resting cage allowing for natural sleep/wake behavior, while being recorded on infrared camera. Sleep were assessed manually based on a 2 min inactivity criteria. Mice received 3x stimulations in each condition (mixed order) in every recording session.

### *In Vivo* Two-photon imaging

16-bit images with double channels and individual frame with spatial dimensions of 512 × 512 was taken at a frequency of 15 Hz. Images were automatically averaged at the rate of 3 to generate 5Hz imaging as output. 920 nm wavelength was applied to capture both GCAMP7 and AlexaFluor594 fluorescence. Agonist Methoxamine (Sigma-Aldrich) was microinjected at a concentration of 50μM into the cortex at the depth of 100 μm using a microelectrode connected to a picospritzer (20 PSI, 20-50 ms; Parker Instrumentation). The microelectrode tip was visualized by adding 100μM AlexaFluor594 (Invitrogen) dissolved in the artificial cerebrospinal fluid (aCSF) pipette solution. Microinjection of 50μM methoxamine was given after 30s recording of spontaneous signals and imaging continued until 100s post-injection. As baseline, two microinjections were given and 15 min of recovery time was allowed between each microinjection. The animal then received i.p. injection of KX or saline. Microinjections at 15 min and 30 min post-injection were given using the same protocol as the baseline condition (one mouse was given KX i.p. injection after 2 microinjections in post-saline state, and another 2 microinjections were conducted at post-KX state). Microelectrodes remained in the same location throughout all imaging sessions in one animal. Efforts were taken to minimize the irritations to the awake mice by avoiding light and noise stimuli during the whole imaging sessions.

Images were analyzed using Fiji. The GCaMP channels were converted to 8-bit and thresholded to capture the calcium signal positive regions properly. “Analyze particles” with particle size setting to be ranging from five pixels to infinity was applied and binary imaging sequences of positive regions for each individual frames were generated. Threshold settings were kept in consistency throughout all the imaging sequences for one animal. Post-injection frames (150-450 frames, post-injection 0-60s) of all the 1^st^ microinjections for each state were selected to make into substacks and the Z-project of 300 frames were generated. Calcium signal propagation regions were subdivided into 1-2 domains according to location of electrode tip and the signal continuity. Maximal propagation area were calculated by manually making a polygon contour inside each domain to cover all the positive regions. Post-injection propagation area was normalized to the pre-injection state for each animal.

### Sleep measurements

Mice were placed in recording chambers (ViewPoint Behavior Technology) and cables were connected to the EEG and EMG electrodes. Cables were connected to a commutator (Plastics One, Bilaney) and the mice were allowed to habituate to the recording chamber (ViewPoint Behavior Technology) for at least one day. On the day of recording, mice were connected to fiber optic implants and recordings were done for 2-4 hours. EEG and EMG signals were amplified (National Instruments Inc.) and filtered (EEG signal: high-pass at 1 Hz and low-pass at 100 Hz; EMG signal: high-pass at 10 Hz and low-pass at 100 Hz), and a notch filter of 50 Hz was used to reduce power line noise. Signals were digitized using a NI Usb 6343 card (National Instrument) and sampled at a sampling rate of 512 Hz. Video was recorded continuously using an infrared video camera (Flir Systems) and used later to aid in the scoring of vigilance states. Hypnograms were created by visual inspection of EEG traces divided into 5 and subsequently one second epochs. Vigilance states were defined as wake (high muscle tonus and a high frequency, low amplitude EEG), nREM sleep (no muscle tonus and low frequency, high amplitude EEG,), and REM sleep (no muscle tonus and high frequency, low amplitude EEG). Analysis of hypnograms was done using SleepScore software (ViewPoint Behavior Technology). All data analysis was subsequently performed in MatLab using custom-made scripts. For a subgroup of animals, sleep measurements were done by activity tracking. Video recordings were done in a white home cage with white bedding and nesting material using an infrared camera. Animals were co-housed with littermates in the white cage at least one day prior to recording, but single-housed during recording. Video recordings were analyzed in Ethovision XT 11.5, in which animals were automatically tracked and sleep was defined as inactivity (activity level < 1.5%) lasting longer than 30 sec. This criteria for sleep was confirmed using combined EEG/EMG and video recording (Suppl. Figure S3).

### Immunohistochemistry

To validate location of optic implant and virus expression, we did immunostaining on brain sections. Animals were deeply anesthetized using KX, then perfused with PBS followed by 4% PFA. Brains were dissected and post-fixed in 4% PFA overnight and transferred to PBS until sectioning. 60 μm sections surrounding the implants were cut using a vibratome. Sections were then blocked in PBS with 5% goat serum and 0.3% Triton X-100 at room temperature for 1-2 hours before overnight incubation with primary antibodies at 4° C. After washing, sections were incubated with secondary antibodies at room temperature for 2 hours, then incubated with DAPI for 2-10 minutes.

Images of whole brain slices were acquired using a Nikon Instruments Ni-E motorized microscope equipped with a 4x CFI Plan Apo Lambda objective (0.2 NA). For excitation, halogen light source were used in combination with excitation filters 362-389 nm, 465-495 nm, 530-575 nm, and Cy5 628640 nm. 4×4 images were acquired per section and stitched together automatically using NIS Elements AR software from Nikon. Close-up images were acquired using a Nikon Instruments C2+ Ti-E Confocal laser-scanning microscope with a 20x CFI Plan Fluor MI objective (0.75 NA) or a 40x CFI Plan Fluor Oil objective (1.30 NA). The excitation sources were 405, 561, and 640 laser diodes and a 488 nm solid-state diode laser. For the 40x images we acquired z-stacks ranging from 20-40 μm and flattened them in Fiji/ImageJ using standard deviation of intensity.

### Primary antibodies

| Name | Company/provider | Cat# |
| --- | --- | --- |
| anti-RFP antibody | Abcam | ab62341 |
| Anti-GFP (Green Fluorescent Protein) (Chicken Antibodies, IgY Fraction) | Aves | GFP-1020 |
| GFP Polyclonal Antibody | Thermo Fisher Scientific | A-6455 |
| anti-Glial Fibrillary Acidic Protein, clone GA5 (mouse monoclonal) | Merck Millipore | MAB360 |
| GFAP Polyclonal Antibody | Invitrogen | PA1-10004 |
| Mouse Anti-Neuronal Nuclei (NeuN) Monoclonal Antibody | Merck Millipore | MAB377 |
| Anti-Tyrosine Hydroxylase, clone LNC1 | Merck Millipore | MAB318 |
| Anti Iba1, Rabbit (for Immunocytochemistry) | Wako | 019-19741 |
| Rabbit Anti-Ng2 Chondroitin Sulfate Proteoglycan Polyclonal Antibody | Merck Millipore | Ab5320 |

### Secondary antibodies

| Name | Company/provider | Cat# |
| --- | --- | --- |
| Goat anti-Rabbit IgG (H+L) Highly Cross-Adsorbed Secondary Antibody, Alexa Fluor 488 | Thermo Fisher Scientific | A11034 |
| Goat anti-Chicken IgY (H+L) Secondary Antibody, Alexa Fluor 488 | Thermo Fisher Scientific | A11039 |
| Goat anti-Rabbit IgG (H+L) Cross-Adsorbed Secondary Antibody, Alexa Fluor 568 | Thermo Fisher Scientific | A11011 |
| Goat anti-Chicken IgY (H+L) Secondary Antibody, Alexa Fluor 568 | Thermo Fisher Scientific | A11041 |
| Goat anti-Mouse IgG1 Cross-Adsorbed Secondary Antibody, Alexa Fluor 647 | Thermo Fisher Scientific | A21240 |

### Statistics

The D’Augustino & Pearson test was used to assess normal distribution of data. The paired t-test or unpaired t-test was employed to compare pairs of groups, if data passed the normality test. Otherwise, the Wilcoxon matched-pairs signed rank test or Mann-Whitney test was used for comparison. For repeated measurements, ANOVA was used.

## Acknowledgements

This work was supported by the Independent Research Council Denmark, the Augustinus Foundation, the Lundbeck Foundation and the Novo Nordisk Foundation.

## Author Contributions

CK: study design, collection of data, analysis, manuscript writing; MA: study design, collection of data, analysis, manuscript writing; NH: EEG surgeries, data collection and analysis; FD: Collect and analyze the calcium data, organize the design and purchase of the NE biosensor virus; WW: supplied the biosensor; QX: Collect two-photon calcium data; SD: supplied the NE biosensor virus, assisted in animal surgery and colony preparation; NK: Collect two-photon calcium data; SP: Collect two-photon calcium data; QS: Optimize and collection of two-photon calcium data; CD: set up and measurements of long-term FP recordings; PKJ: constructed and provided GFAP-virus; JF: constructed and provided NE biosensor virus; YL: constructed and provided NE biosensor virus; PW: study design and data discussion; HH: study design, data organizing and discussion; MN: study design, data organizing and discussion.

## Reference list

Agster, K.L., Mejias-Aponte, C.A., Clark, B.D., and Waterhouse, B.D. (2013). Evidence for a Regional Specificity in the Density and Distribution of Noradrenergic Varicosities in Rat Cortex. J. Comp. Neurol. 521.

Andrillon, T., Nir, Y., Staba, R.J., Ferrarelli, F., Cirelli, C., Tononi, G., and Fried, I. (2011). Sleep spindles in humans: insights from intracranial EEG and unit recordings. J. Neurosci. 31, 17821–17834.

Aston-Jones, G., and Bloom, F.E. (1981). Activity of norepinephrine-containing locus coeruleus neurons in behaving rats anticipates fluctuations in the sleep-waking cycle. J. Neurosci. 1, 876–886.

Aston-Jones, G., and Cohen, J.D. (2005). An integrative theory of locus coeruleus-norepinephrine function: adaptive gain and optimal performance. Annu. Rev. Neurosci. 28, 403–450.

Bekar, L.K., He, W., and Nedergaard, M. (2008). Locus coeruleus alpha-adrenergic-mediated activation of cortical astrocytes in vivo. Cereb. Cortex 18, 2789–2795.

Bellesi, M., Tononi, G., Cirelli, C., and Serra, P.A. (2016). Region-Specific Dissociation between Cortical Noradrenaline Levels and the Sleep/Wake Cycle. Sleep 39, 143–154.

Berridge, C.W., and Abercrombie, E.D. (1999). Relationship between locus coeruleus discharge rates and rates of norepinephrine release within neocortex as assessed by in vivo microdialysis. Neuroscience 93, 1263–1270.

Berridge, C.W., Page, M.E., Valentino, R.J., and Foote, S.L. (1993). Effects of locus coeruleus inactivation on electroencephalographic activity in neocortex and hippocampus. Neuroscience 55, 381–393.

Bojarskaite, L., Bjørnstad, D.M., Pettersen, K.H., Cunen, C., Hermansen, G.H., Åbjørsbråten, K.S., Chambers, A.R., Sprengel, R., Vervaeke, K., Tang, W., et al. (2020). Astrocytic Ca2+ signaling is reduced during sleep and is involved in the regulation of slow wave sleep. Nat. Commun. 11, 3240.

Brown, R.E., Basheer, R., McKenna, J.T., Strecker, R.E., and McCarley, R.W. (2012). Control of Sleep and Wakefulness. Physiol. Rev. 92, 1087–1187.

Cao, X., Li, L.-P., Wang, Q., Wu, Q., Hu, H.-H., Zhang, M., Fang, Y.-Y., Zhang, J., Li, S.-J., Xiong, W.-C., et al. (2013). Astrocyte-derived ATP modulates depressive-like behaviors. Nat. Med. 19, 773–777.

Carter, M.E., Yizhar, O., Chikahisa, S., Nguyen, H., Adamantidis, A., Nishino, S., Deisseroth, K., and de Lecea, L. (2010). Tuning arousal with optogenetic modulation of locus coeruleus neurons. Nat. Neurosci. 13, 1526–1533.

Chandler, D.J., Gao, W.-J., and Waterhouse, B.D. (2014). Heterogeneous organization of the locus coeruleus projections to prefrontal and motor cortices. Proc. Natl. Acad. Sci. U. S. A. 111, 6816–6821.

Ding, F., O’Donnell, J., Thrane, A.S., Zeppenfeld, D., Kang, H., Xie, L., Wang, F., and Nedergaard, M. (2013). α1-Adrenergic receptors mediate coordinated Ca2+ signaling of cortical astrocytes in awake, behaving mice. Cell Calcium 54, 387–394.

Ding, F., O’Donnell, J., Xu, Q., Kang, N., Goldman, N., and Nedergaard, M. (2016). Changes in the composition of brain interstitial ions control the sleep-wake cycle. Science 352, 550–555.

Eschenko, O., and Sara, S.J. (2008). Learning-dependent, transient increase of activity in noradrenergic neurons of locus coeruleus during slow wave sleep in the rat: brain stem-cortex interplay for memory consolidation? Cereb. Cortex 18, 2596–2603.

Eschenko, O., Magri, C., Panzeri, S., and Sara, S.J. (2012). Noradrenergic Neurons of the Locus Coeruleus Are Phase Locked to Cortical Up-Down States during Sleep. Cereb. Cortex 22, 426–435.

Feng, J., Zhang, C., Lischinsky, J.E., Jing, M., Zhou, J., Wang, H., Zhang, Y., Dong, A., Wu, Z., Wu, H., et al. (2019). A Genetically Encoded Fluorescent Sensor for Rapid and Specific In Vivo Detection of Norepinephrine. Neuron 102, 745–761.

Florin-Lechner, S.M., Druhan, J.P., Aston-Jones, G., and Valentino, R.J. (1996). Enhanced norepinephrine release in prefrontal cortex with burst stimulation of the locus coeruleus. Brain Res. 742, 89–97.

Foley, J., Blutstein, T., Lee, S., Erneux, C., Halassa, M.M., and Haydon, P. (2017). Astrocytic IP3/Ca2+ Signaling Modulates Theta Rhythm and REM Sleep. Front. Neural Circuits 11, 3.

Foote, S.L., Aston-Jones, G., and Bloom, F.E. (1980). Impulse activity of locus coeruleus neurons in awake rats and monkeys is a function of sensory stimulation and arousal. Proc. Natl. Acad. Sci. U. S. A. 77, 3033–3037.

Gervasoni, D., Panconi, E., Henninot, V., Boissard, R., Barbagli, B., Fort, P., and Luppi, P.. (2002). Effect of chronic treatment with milnacipran on sleep architecture in rats compared with paroxetine and imipramine. Pharmacol. Biochem. Behav. 73, 557–563.

Hayat, H., Regev, N., Matosevich, N., Sales, A., Paredes-Rodriguez, E., Krom, A.J., Bergman, L., Li, Y., Lavigne, M., Kremer, E.J., et al. (2020). Locus coeruleus norepinephrine activity mediates sensory-evoked awakenings from sleep. Sci. Adv. 6.

Hobson, J.A., McCarley, R.W., and Wyzinski, P.W. (1975). Sleep cycle oscillation: reciprocal discharge by two brainstem neuronal groups. Science 189, 55–58.

Hunsley, M.S., and Palmiter, R.D. (2004). Altered sleep latency and arousal regulation in mice lacking norepinephrine. Pharmacol. Biochem. Behav. 78, 765–773.

Ingiosi, A.M., Hayworth, C.R., Harvey, D.O., Singletary, K.G., Rempe, M.J., Wisor, J.P., and Frank, M.G. (2019). A role for astroglial calcium in mammalian sleep. BioRxiv 728931.

Jones, B.E. (1991). The role of noradrenergic locus coeruleus neurons and neighboring cholinergic neurons of the pontomesencephalic tegmentum in sleep-wake states. Prog. Brain Res. 88, 533–543.

Lee, K.H., and McCormick, D.A. (1996). Abolition of Spindle Oscillations by Serotonin and Norepinephrine in the Ferret Lateral Geniculate and Perigeniculate Nuclei In Vitro. Neuron 17, 309–321.

Léna, I., Parrot, S., Deschaux, O., Muffat-Joly, S., Sauvinet, V., Renaud, B., Suaud-Chagny, M.-F., and Gottesmann, C. (2005). Variations in extracellular levels of dopamine, noradrenaline, glutamate, and aspartate across the sleep-wake cycle in the medial prefrontal cortex and nucleus accumbens of freely moving rats. J. Neurosci. Res. 81, 891–899.

Li, W., Ma, L., Yang, G., and Gan, W.-B. (2017). REM sleep selectively prunes and maintains new synapses in development and learning. Nat. Neurosci. 20, 427–437.

Li, Y., Hickey, L., Perrins, R., Werlen, E., Patel, A.A., Hirschberg, S., Jones, M.W., Salinas, S., Kremer, E.J., and Pickering, A.E. (2016). Retrograde optogenetic characterization of the pontospinal module of the locus coeruleus with a canine adenoviral vector. Brain Res. 1641, 274.

Monai, H., Ohkura, M., Tanaka, M., Oe, Y., Konno, A., Hirai, H., Mikoshiba, K., Itohara, S., Nakai, J., Iwai, Y., et al. (2016). Calcium imaging reveals glial involvement in transcranial direct current stimulation-induced plasticity in mouse brain. Nat. Commun. 7, 11100.

Monti, J.M., D’Angelo, L., Jantos, H., Barbeito, L., and Abó, V. (1988). Effect of DSP-4, a noradrenergic neurotoxin, on sleep and wakefulness and sensitivity to drugs acting on adrenergic receptors in the rat. Sleep 11, 370–377.

Oe, Y., Wang, X., Patriarchi, T., Konno, A., Ozawa, K., Yahagi, K., Hirai, H., Tsuboi, T., Kitaguchi, T., Tian, L., et al. (2020). Distinct temporal integration of noradrenaline signaling by astrocytic second messengers during vigilance. Nat. Commun. 11, 471.

Paukert, M., Agarwal, A., Cha, J., Doze, V.A., Kang, J.U., and Bergles, D.E. (2014). Norepinephrine controls astroglial responsiveness to local circuit activity. Neuron 82, 1263–1270.

Porter-Stransky, K.A., Centanni, S.W., Karne, S.L., Odil, L.M., Fekir, S., Wong, J.C., Jerome, C., Mitchell, H.A., Escayg, A., Pedersen, N.P., et al. (2019). Noradrenergic Transmission at Alpha1-Adrenergic Receptors in the Ventral Periaqueductal Gray Modulates Arousal. Biol. Psychiatry 85, 237–247.

Rajkowski, J., Kubiak, P., and Aston-Jones, G. (1994). Locus coeruleus activity in monkey: Phasic and tonic changes are associated with altered vigilance. Brain Res. Bull. 35, 607–616.

Rasmussen, K., Morilak, D.A., and Jacobs, B.L. (1986). Single unit activity of locus coeruleus neurons in the freely moving cat. I. During naturalistic behaviors and in response to simple and complex stimuli. Brain Res. 371, 324–334.

Sardinha, V.M., Guerra-Gomes, S., Caetano, I., Tavares, G., Martins, M., Reis, J.S., Correia, J.S., Teixeira-Castro, A., Pinto, L., Sousa, N., et al. (2017). Astrocytic signaling supports hippocampal-prefrontal theta synchronization and cognitive function. Glia 65, 1944–1960.

Seibt, J., Richard, C.J., Sigl-Glöckner, J., Takahashi, N., Kaplan, D.I., Doron, G., de Limoges, D., Bocklisch, C., and Larkum, M.E. (2017). Cortical dendritic activity correlates with spindle-rich oscillations during sleep in rodents. Nat. Commun. 8, 684.

Shouse, M.N., Staba, R.J., Saquib, S.F., and Farber, P.R. (2000). Monoamines and sleep: microdialysis findings in pons and amygdala. Brain Res. 860, 181–189.

Singh, S., and Mallick, B.N. (1996). Mild electrical stimulation of pontine tegmentum around locus coeruleus reduces rapid eye movement sleep in rats. Neurosci. Res. 24, 227–235.

Steriade, M., Timofeev, I., and Grenier, F. (2001). Natural Waking and Sleep States: A View From Inside Neocortical Neurons. J. Neurophysiol. 85, 1969–1985.

Swift, K.M., Gross, B.A., Frazer, M.A., Bauer, D.S., Clark, K.J.D., Vazey, E.M., Aston-Jones, G., Li, Y., Pickering, A.E., Sara, S.J., et al. (2018). Abnormal Locus Coeruleus Sleep Activity Alters Sleep Signatures of Memory Consolidation and Impairs Place Cell Stability and Spatial Memory. Curr. Biol. 28, 3599–3609.

Takahashi, K., Kayama, Y., Lin, J.S., and Sakai, K. (2010). Locus coeruleus neuronal activity during the sleep-waking cycle in mice. Neuroscience 169, 1115–1126.

Thrane, A.S., Rangroo Thrane, V., Zeppenfeld, D., Lou, N., Xu, Q., Nagelhus, E.A., and Nedergaard, M. (2012). General anesthesia selectively disrupts astrocyte calcium signaling in the awake mouse cortex. Proc. Natl. Acad. Sci. U. S. A. 109, 18974–18979.

Tian, G.-F., Azmi, H., Takano, T., Xu, Q., Peng, W., Lin, J., Oberheim, N., Lou, N., Wang, X., Zielke, H.R., et al. (2005). An astrocytic basis of epilepsy. Nat. Med. 11, 973–981.

Ulrich, D. (2016). Sleep Spindles as Facilitators of Memory Formation and Learning. Neural Plast. 2016, 1–7.

Yamaguchi, H., Hopf, F.W., Li, S.-B., and Lecea, L. de (2018). In vivo cell type-specific CRISPR knockdown of dopamine beta hydroxylase reduces locus coeruleus evoked wakefulness. Nat. Commun. 9, 5211.

