## Supplementary data for "Dynamic fluctuations of the locus coeruleus-norepinephrine system underlie sleep state transitions"

The supplementary data contains:

Supplementary Figure S1 (related to Figure 1)

Supplementary Figure S2 (related to Figure 1)

Supplementary Figure S3 (related to Figure 1)

Supplementary Figure S4 (related to Figure 1)

Supplementary Figure S5 (related to Figure 2)

Supplementary Figure S6 (related to Figure 3)

Supplementary Figure S7 (related to Figure 3)

**Figure S1**

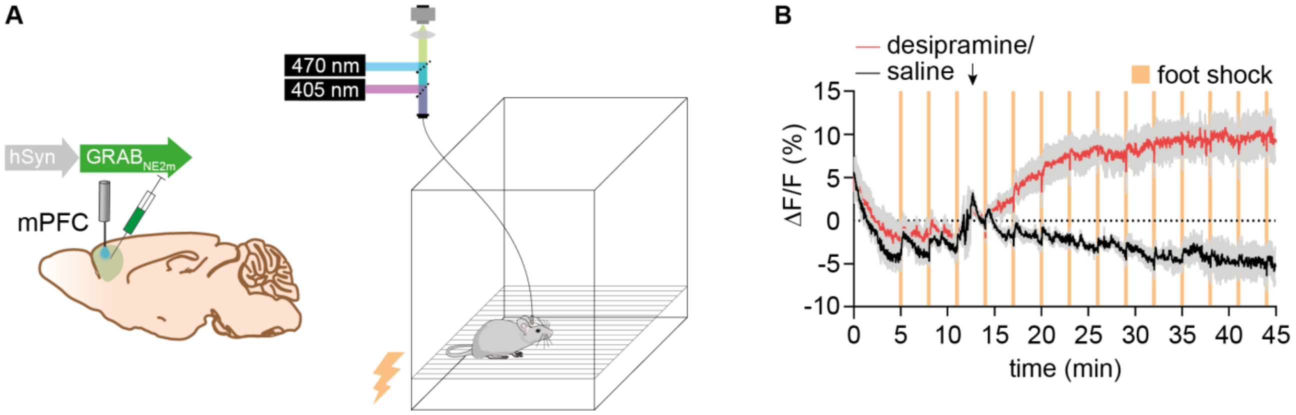


**Supplementary Figure S1. Inhibition of norepinephrine (NE) reuptake increases NE biosensor signal**. **A.** GRAB$_{NE2m}$ was expressed in mPFC under the neuronal hSyn promoter and mice were subjected to repeated foot shocks (0.5 s 0.45 mA with 3 min intervals) while we recorded GRAB$_{NE2m}$ fluorescence by fiber photometry. **B.** After the first three foot shocks, mice received a saline or desipramine (10 mg/kg, i.p.) injection and the mean traces of the two groups are shown here. Data is shown as mean±SEM, n = 6.

**Figure S2**

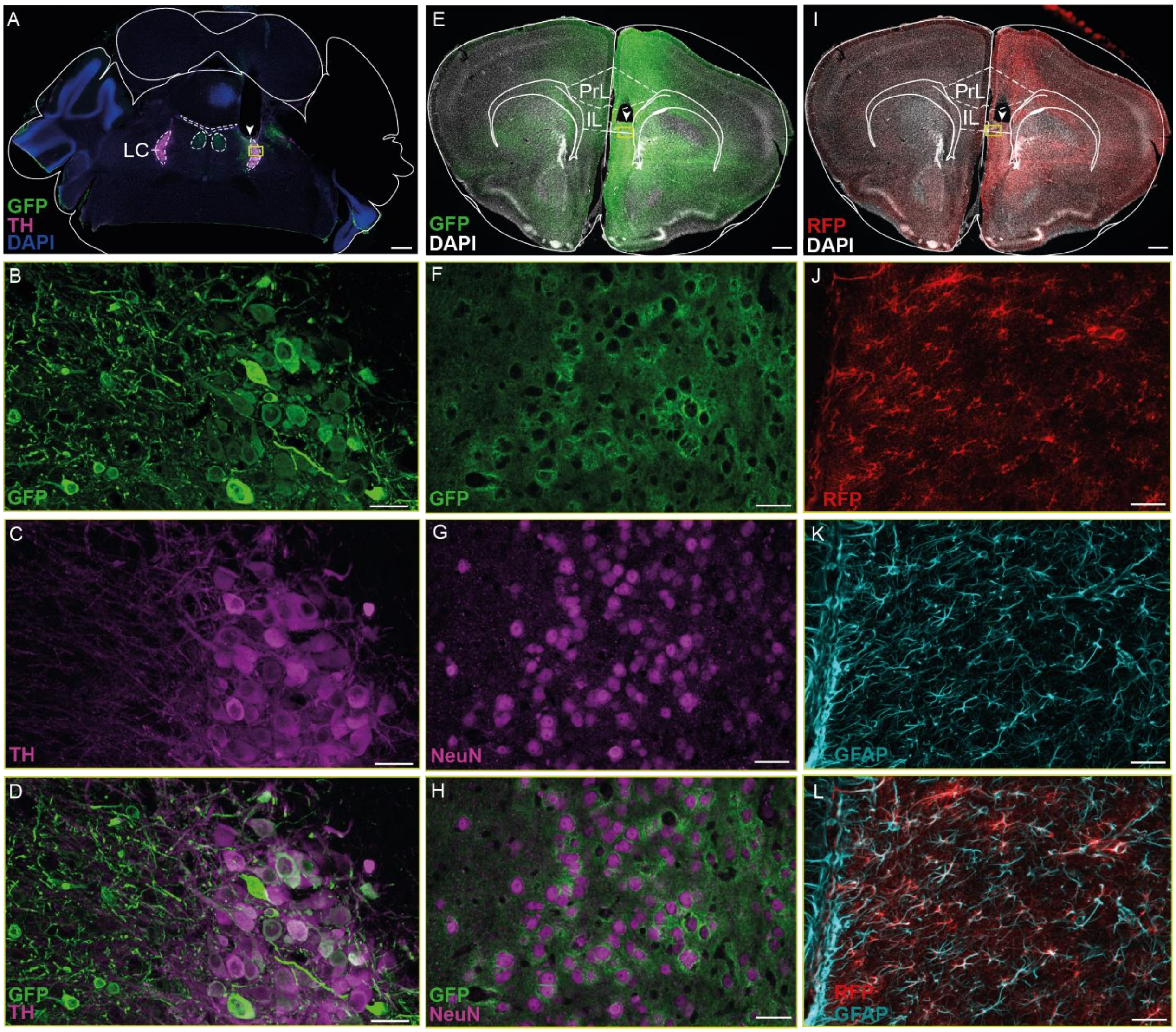


**Supplementary Figure S2. Verification of virus expression and optical fiber location. A-D.** Expression of GCaMP6f in tyrosine hydroxylase (TH) positive neurons of LC was verified by co-staining for TH and green fluorescent protein (GFP). **E-H.** Expression of the norepinephrine biosensor, $GRAB_{NE2m}$, in neurons of medial prefrontal cortex (mPFC) was verified by co-staining for GFP and neuronal nuclei (NeuN). **I-L**. Astrocytic expression of jRGECO1a in mPFC was verified by co-staining of glial fibrillary acidic protein (GFAP) and red fluorescent protein (RFP). Scale bars are 400 µm (**A**, **E** and **I**), and 30 µm (**B-D**, **F-H**, and **J-K**). Arrowheads mark the site of the optical fiber tip and yellow squares shows location of magnified images.

**Figure S3**

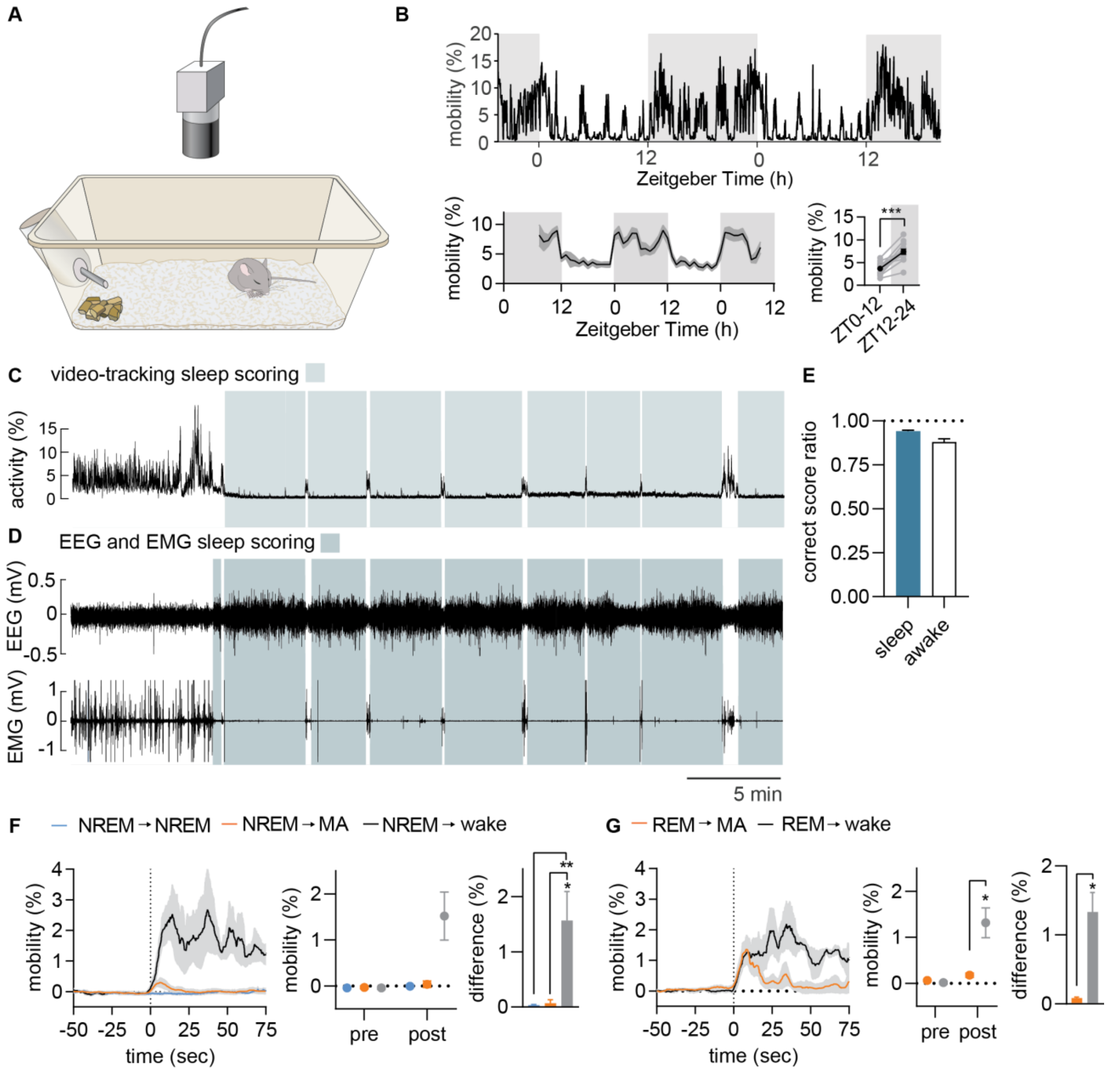


**Supplementary Figure S3. Mobilization status of the mouse marks sleep to wake transitions.** **A.** The mobility of the mouse was tracked using videos recorded by a camera placed over a cage containing white bedding. **B.** At the top is shown example data of mobility over the course of 48 hours. At the lower panel is displayed summary data (grey zone: dark phase; white zone: light phase). Mice move more during the dark than the light phase **C.** Example data of mobility performed by videotracking of a mouse implanted with EEG and EMG electrodes. Green zones indicate video-based sleep scoring (30 sec of <1.5% mobility) **D.** Example data of EEG and EMG recordings corresponding to the video-based mobility data above. Green zones indicate EEG and

EMG based sleep scoring. **E.** Summary plot of the percentage of video-based sleep scorings that was correct (compared to EEG/EMG scoring). **F.** In the left panel is the mean video-based mobility across NREM to NREM, NREM to microarousal (MA) and NREM to wake transitions. The middle panel shows mean mobility values pre and post transition and the right panel shows the difference (post minus pre) for the different transitions. **G.** Similar to **F**, but for REM to microarousal (MA) and REM to wake transitions. Paired t test, n=12 **(A-E)**, n = 6 **(F-G)**. Data is shown as mean±SEM. *$p < 0.05$, **$p < 0.01$, ***$p < 0.001$.

**Figure S4**

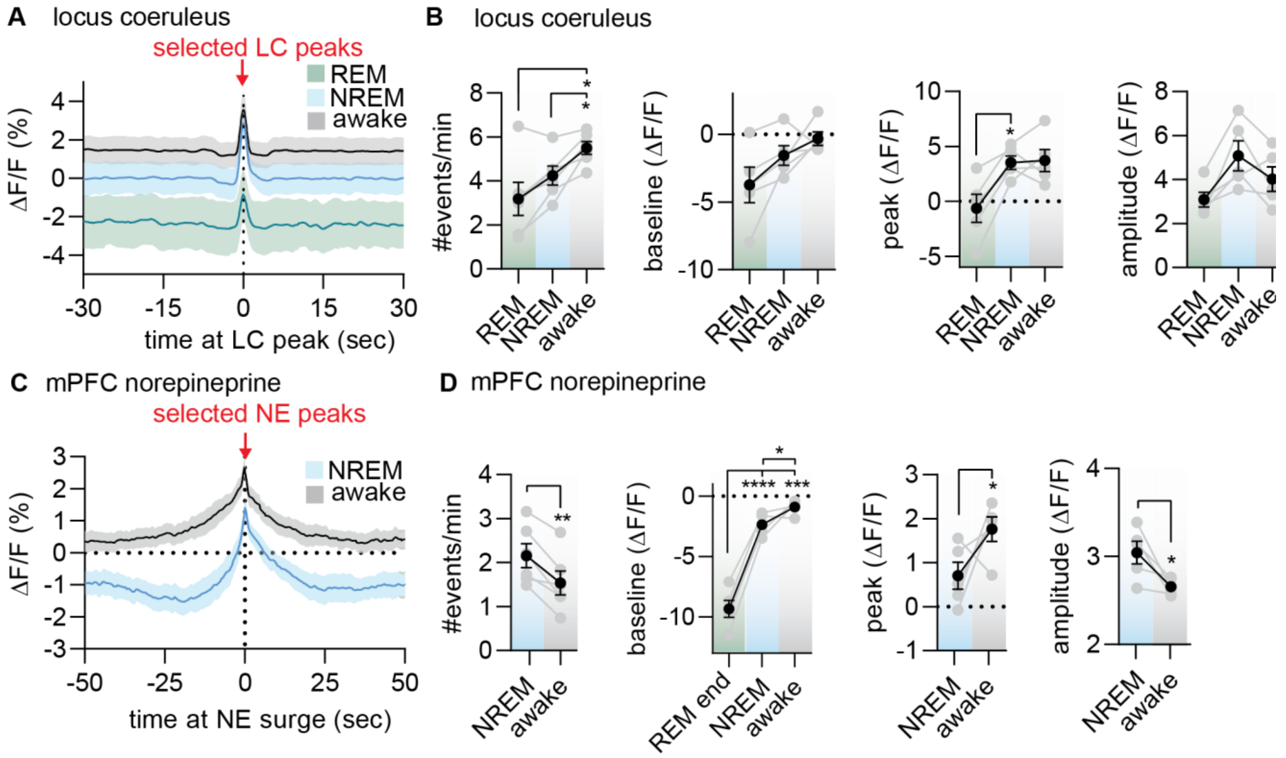


**Supplementary Figure S4. Sleep state dependent locus coeruleus-norepinephrine activity. A.** Locus coeruleus (LC) peaks were selected and categorized into the brain state they occurred in: wakefulness, NREM or REM. **B.** Summary of the frequency and kinetics of the selected LC peaks dependent on brain state. There are most LC events during wake followed by NREM and REM. The selected peak values during REM are smaller than NREM and wake and might be attributed to noise. **C.** The peaks of norepinephrine (NE) increases were selected and the events were divided into sleep states (no NE peaks were apparent during REM). **D.** Summary plots showing the frequency and kinetics of the NE events. There are more NE event during NREM compared to wake probably due to the larger dynamic NE range during sleep, which is also reflected in the larger peak and smaller amplitude of wake-related NE events. The NE baseline value was largest during awake, and lowest of the end of REM. Data is shown as mean±SEM, n = 5-6, *p < 0.05, **p < 0.01, ***p < 0.001, ****p < 0.0001.

**Figure S5**

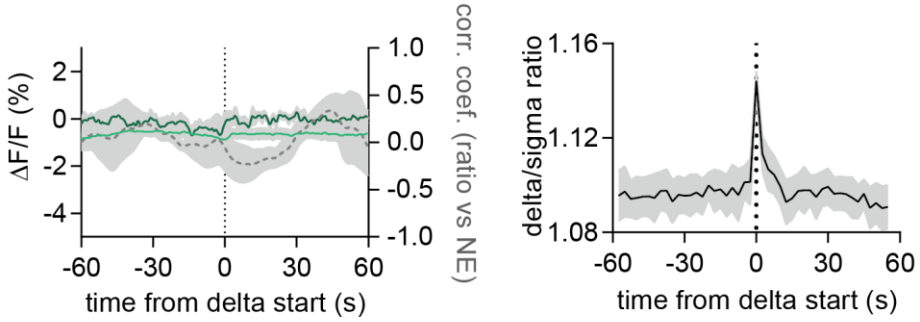


**Supplementary Figure S5. Delta activity does not correlate with norepinephrine levels.** Left: Mean LC (dark green) and NE (green) traces aligned to time of high delta activity during NREM. The dotted grey trace is the correlation coefficient between delta/sigma ratio (shown in right part of figure) and NE level. Data is shown as mean±SEM, n = 5.

**Figure S6**

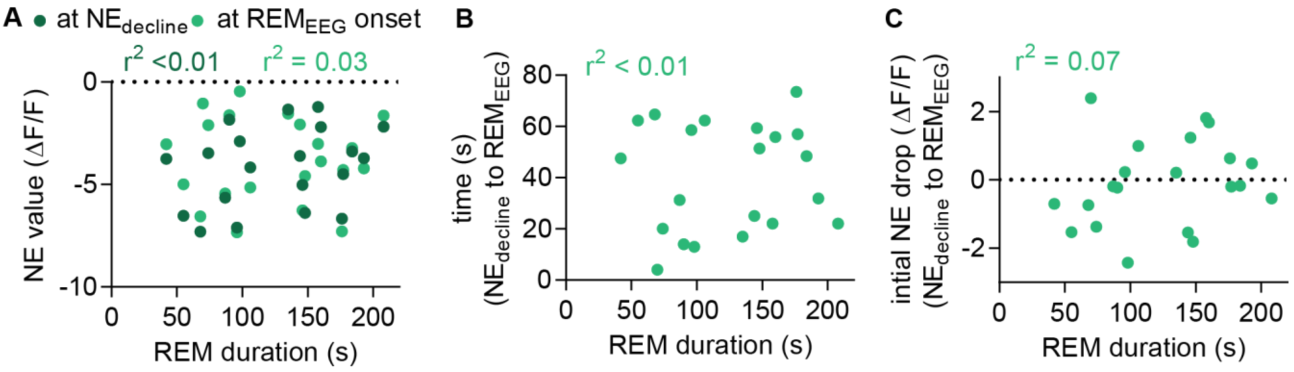


**Supplementary Figure S6. Norepinephrine level preceding REM does not correlate with REM duration. A.** There was no correlation between REM duration and the level of norepinephrine (NE) prior to (dark green) or at REM onset (light green). **B.** There was no correlation between REM duration and the time between onset of NE decline and onset of REM onset. **C.** There was no correlation between REM duration and the NE drop between onset of NE decline and REM onset (C). Data is shown as mean±SEM, n = 6.

### Figure S7

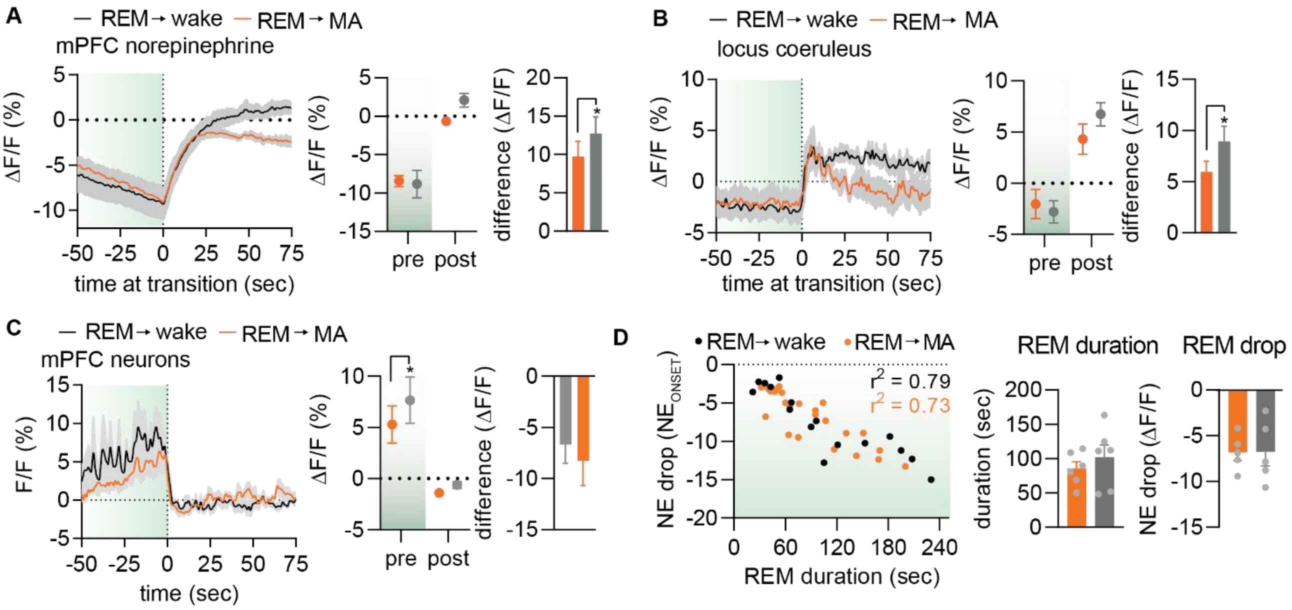


**Supplementary Figure S7. The impact of different behavioral outcomes at REM offset.** Mean norepinephrine (NE) (**A**), locus coeruleus (LC) (**B**), and medial prefrontal (mPFC) neuronal (**C**) traces aligned to the point of NE increase preceding REM offset. Black/grey illustrates transitions to wakefulness and orange to microarousals (MA). Middle panels show mean levels prior to (-10 to 0 s, 'pre') and after (max value, 0-50 s, 'post') transition and right panels show difference between 'post' and 'pre' levels. **D.** Correlation between REM duration and NE drop (left), mean REM duration (middle) and mean NE drop (right) categorized into REM bouts followed by wakening or MA. Data is shown as mean±SEM, n = 6, *$p < 0.05$, **$p < 0.01$, ***$p < 0.001$, ****$p < 0.0001$.
